# Sick and Tired: Polyparasitism, Food Stress and Sickness Behaviour in a Gregarious Mammal

**DOI:** 10.1101/2021.04.22.440985

**Authors:** Rosemary Blersch, Tyler R. Bonnell, Andre Ganswindt, Christopher Young, Louise Barrett, S. Peter Henzi

## Abstract

Although sickness behaviour in response to non-lethal parasites has been documented in wild animals, it remains unclear how social and environmental stress might also shape an animal’s behavioural response to parasitism, nor do we know whether simultaneous infection with more than one parasite changes the way animals respond. Here, we combine physiological, environmental, behavioural and parasite measures to investigate behavioural responses to infection in wild vervet monkeys (Chlorocebus pygerythrus) living in a semi-arid region of South Africa. We quantified both activity budget and behavioural predictability to investigate the occurrence of sickness behaviour and infection with two non-lethal gastrointestinal parasite genera. Higher parasite load was linked to an increase in the time spent resting. However, the nature of the relationship with other behaviours was contingent on both the parasite genus in question, and how parasite species interacted, highlighting the importance of considering co-infection. Overall, food availability was the dominant predictor of behavioural change suggesting that, for monkeys living in a more extreme environment, coping with ecological stress may override the ability to modulate behaviour in response to other physiological stressors. Our findings provide insight into how animals living in harsh environments find ways to cope with parasite infection, avoidance, and transmission.

## Introduction

It has long been established that pathogens with high fatality rates, such as epidemic viruses, can drive population declines, and may contribute to local extinctions [1]. The effects of sub-clinical or non-lethal infections are often overlooked, however, despite costs to host health and fitness, and the consequent impact of such diseases on population viability [2]. Hosts have evolved several physiological and behavioural responses to cope with the pressures of infection [3] and, while we have a firm grasp on the physiological immune response to infections in wildlife, less is known about the behavioural presentation of sickness and its physiological correlates [4].

Sickness behaviour is broadly defined as a suite of behaviours that occurs in response to infection. This includes lethargy, anorexia, somnolence, and a reduction in grooming [4,5]. Although originally thought to be simply a by-product of illness, sickness behaviour is now considered to be part of a highly organised strategy to combat infection by reallocating energy to the immune system and away from non-essential activities [5]. While exhibiting sickness behaviour is inherently beneficial to combatting infection, a trade-off emerges as energetic resources are devoted to fighting infection at the expense of other vital processes, such as growth and reproduction [3]. The severity of these costs, and hence the relative benefit of displaying sickness behaviour, depends on ecological context and the value of behaviours that need to be sacrificed. Thus, we should expect to see animals modulating their expression of sickness behaviours when the costs become too high; something particularly pertinent to animals subject to prolonged environmental or social stress [6,7].

Sickness behaviour has been extensively documented in captive populations [2,8–10], but we know much less about its occurrence in wild mammals [11–13]—most likely due to the challenges associated with long-term environmental and physiological monitoring. Sickness behaviour research in the wild, therefore, has focused almost exclusively on the relationship between parasite infection and behaviour, independent of other stressors.

However, the reorganisation of behaviour is more complex if animals are simultaneously subject to other competing stressors common in natural environments [6,7], and the expression of sickness behaviour should vary accordingly. Although we have some grasp of the social factors that influence investment in sickness behaviour (for review see [3]), the influence of environmental stressors remains poorly understood. Understanding the interplay between environmental stress and behavioural modification is central to understanding how sickness behaviour may impact long-term fitness in wild populations.

Sickness behaviour research has also been principally concerned with the effects of a single designated parasite or pathogen species on behaviour. Yet, wild animals rarely harbour only a single species, and interactions between parasite species are likely [14]. This interaction can be either synergic, where the parasite burden of one species magnifies the consequences of another, or antagonistic, where the burden supresses the other’s effects [15]. At present, we have evidence that polyparasitism predicts infection risk [16], host body condition, and survival [17] in mammals but we there is comparatively little research on how multi-parasite infection affects behaviour (see: [18–23]).

Here, we use a comprehensive dataset comprised of detailed physiological (faecal glucocorticoid metabolites), environmental, behavioural and parasite data to assess how these factors interact to shape behavioural responses to infection in a population of a highly social, wild mammal, specifically, the vervet monkey (*Chlorocebus pygerythrus*), in a semi-arid region of South Africa. Previous work in this population has identified complex relationships between behaviour and environmental conditions, with food resources, temperature, rainfall, and standing water availability strongly influencing activity budgets and mortality [25,26]. As in this previous work, we use fGCMs as an index of individual response to environmental stressors (i.e., as a measure of the ability to restore homeostasis), rather than an indicator of an individual animals stress levels [24]. Given the often harsh environmental conditions in the study area, these monkeys provide an excellent opportunity to determine whether the expression of sickness behaviour in wild animals is modulated, or suppressed, by other simultaneous external and internal stressors.

We use a combined approach, quantifying both activity budget and behavioural predictability, to investigate the relationships between behaviour and two non-lethal gastrointestinal parasite genera. In addition to a more comprehensive dataset, we use a newly developed measure of entropy rate to assess predictability [27]; this allows a larger range of behaviours to be considered, and is therefore more sensitive than existing analytical techniques. Finally, we consider whether there is an interaction between the two parasite genera studied here, and if co-infection compounds the need to invest in sickness behaviours.

## Methods

### Study Site and Study Species

We collected data and faecal samples from August 2017 to April 2018 from three fully habituated groups (PT = Picnic Troop, RBM = River Bend Mob, RST = Riverside Troop) of wild vervet monkeys on Samara Private Game Reserve, South Africa (32°22’S, 24°52’E). These monkeys have been the subject of continuous data collection since 2009. All group members were individually identified based on natural markings, and data for this study were collected from a subset of 27 adult individuals (PT: 4 males, 6 females out of 14 adults; RBM: 2 males, 6 females out of 14 adults; RST: 3 males, 6 females out of 16 adults), selected to be representative of adult demography and to reflect the full range of dominance ranks. The study area comprises semi-arid riverine woodland [28], with a declining annual average rainfall of 386 mm, and average annual minimum and maximum temperatures of 10°C and 27°C respectively. The region experiences periodic droughts that are severe enough to be a primary source of mortality for animals in our study groups [26].

### Behavioural Data Collection

Each group was followed for five days each week across the study period, and data were collected for 10 hours each day [26,29]. To assess changes in activity budget, the behaviour of all visible individuals was recorded during 10-min scan sampling blocks [30] conducted every 30 min throughout the day. We selected four, high frequency, mutually exclusive behaviours for analysis: moving, foraging, resting and allo-grooming, either given or received. Notably, we considered foraging to include both manipulation and ingestion of food (see [31] for definitions).

To investigate changes in behavioural predictability, we conducted 10-min continuous focal sampling [30] twice per week for each of the 27 subjects (N_total_ =1614 focal samples). Randomised focal times were generated for each day. During these focal sampling events, a single individual was followed and a continuous, timed record of its behaviour obtained, using electronic data loggers and proprietary software. The same mutually exclusive behaviours were identified as described above. Owing either to disruptions, such as aggressive encounters between groups, or periods where individuals were out of sight, not all focal samples were exactly 10 minutes long. We selected 6 min as the cut-off time for inclusion in the data set and controlled for focal sample length in our analyses.

Finally, we collected *ad libitum* data on dyadic agonistic interactions among all group members, for which we identified participants and outcomes. Given good visibility at the site we are confident that there was no systematic bias in the likelihood of observing encounters. These agonistic data were used to construct dominance hierarchies [26]. Only decided dyadic agonistic interactions with a clear winner and loser were included in the analysis.

### Dominance Hierarchy

We divided the study period into four 3-month blocks: July – September 2017, October – December 2017, January – March 2018 and April – June 2018. We used *ad libitum* observations of agonistic interactions to construct hierarchies for each period (RBM_Total N_: 963; RST_Total N_: 810; PT_Total N_: 1135) for all adults in each troop and not only the subset of study subjects. Given male-female co-dominance in this population [32], we generated a single matrix that included all decided agonistic interactions regardless of the sex of participants and created a single interdigitated hierarchy.

Dominance ranks in each troop and for each 3-month block were expressed as a standardized David’s score using the package ‘compete’ [33]. David’s scores were standardized to enable direct comparison across groups of different size and interaction rates [34].

### Food availability

We quantified food availability in each troop’s home range by calculating the Normalized Difference Vegetation Index (NDVI) every 16 days [26] from MODIS data collected by Earth Observing System (EOS) satellites Terra (EOS AM-1) and Aqua (EOS PM-1). Using Moderate Resolution Imaging Spectroradiometer MOD13Q1 vegetation indices at a 250-meter resolution [35], NDVI measures the amount of biomass or chlorophyll activity by calculating the difference between the visible red and near infrared bands divided by their sum. The resultant measure is a range of values between −1 and 1, where negative values indicate an absence of vegetation and positive values approaching 1 indicate larger concentrations of green vegetation [36]. Given the generalist, largely plant-based nature of vervet diet [28], the synoptic view of NDVI is a reliable measure of food availability in this species and at this site [37,38].

### Faecal sampling and analysis

We collected a total of 573 faecal samples (mean = 21/individual, ± 3.1 SD) during the 234 days of the study. Faecal samples were collected twice per month (once during each two-week period) from the 27 subjects. Two corresponding faecal samples, one for parasite analysis and one for faecal glucocorticoid metabolites (fGCM) analysis, were collected from the same defecation event.

### Parasite analysis

For each sample, approximately 1 g of fresh faeces was weighed in the field immediately after defecation and directly placed into 10% neutral, buffered formalin. Samples were stored in the field lab and transported to the University of Lethbridge, Canada, where faecal flotation and sedimentation techniques were used to identify parasites [39].

We used a modified zinc sulphate flotation to isolate helminth eggs followed by ethyl-acetate sedimentation to isolate potential trematodes that were too heavy to float during ZnSO4 flotation (methods: supplementary S1). For both methods, the entire pellet was examined under the microscope. Parasites were identified to genus- level based on egg shape, size, colour, and contents, and all eggs were counted. Representative eggs were photographed.

We recovered 5 parasite genera from faecal samples [39]. One parasite could not be identified to genus level, as eggs of *Physaloptera* sp. and *Protospirura* sp. cannot be reliably distinguished based on egg morphology alone. Based on morphological characteristics of the eggs, including their size and the presence of a hyaline substance [40,41], we consider it most likely to be *Protospirura* sp. (hereafter referred to as ?*Protospirura* sp.) pending results of ongoing molecular analysis. Due to small sample size for three genera (<5% mean annual sample prevalence), namely *Oesophagostomum* sp., *Subulura* sp. and *Ternidens* sp., we selected only ?*Protospirura* sp. and *Trichostrongylus* sp. (>20% mean annual sample prevalence) for these analyses but include other species in the number of genera (parasite richness).

We have established previously that sequential faecal egg count patterns for *Trichostrongylus* sp. and ?*Protospirura* sp. are not stochastic and point to underlying levels of infection in our population [37, Blersch et al. in press], and thus use egg counts as a proxy for the extent of helminth infection.

### Faecal steroid analysis

Samples were collected following the protocol of Young et al. [26,42]. Within 15min of defecation, a 2-5g piece of faecal material was transferred into a plastic vial following physical homogenization of the full faecal sample. Prior to collection, faecal samples were checked to ensure there was no contamination with urine during excretion or on the substrate where the sample landed. Vials were immediately stored on ice in a thermos flask in the field before transfer to a −20°C freezer at the end of the day. Samples were stored frozen until transport on dry ice to the Endocrine Research Laboratory, University of Pretoria, for analysis.

Samples were lyophilized, pulverized and then sieved to remove seeds and fibrous matter [42]. The resulting faecal powder (0.10g) was extracted by vortexing for 15min with 80% ethanol in water (3ml) followed by 10 minutes of centrifugation at 1500g. 1.5 ml of the resultant supernatants were transferred into microcentrifuge tubes. Hormone analysis was conducted following the standard procedures of the Endocrine Research Laboratory, University of Pretoria [43] using the cortisol enzyme immunoassay (EIA) [42]. Seinsitivity of the EIA was 0.6 ng/g dry weight [42]. Inter- and intra-assay coefficients of variation of high- and low-value quality controls were: 4.64–5.96 and 8.13–11.60% respectively. All steroid concentrations are given as ng g^-1^ faecal dry weight.

## Statistical Analysis

### Patterns of co-infection

Egg counts of our two most prevalent parasite genera, ?*Protospirura* sp. and *Trichostrongylus* sp., were used in these analyses. We conducted exploratory analysis to assess whether there was a relationship in parasite intensity between ?*Protospirura* sp. and *Trichostrongylus* sp., using a mixed effects model in a Bayesian framework and specifying a lognormal distribution. We filtered out samples that were parasite negative (N = 8). ?*Protospirura* sp. intensity, represented as eggs per gram (EPG) was our response variable while *Trichostrongylus* sp. was our fixed effect. We included individual ID nested in troop as our random effect with individual-level random slopes for *Trichostrongylus* sp.

### Model set 1: The influence of parasite infection and ecology on behaviour

To examine whether infection with ?*Protospirura* sp., *Trichostrongylus* sp. and parasite species richness (the number of genera recovered in each faecal sample) were associated with changes in behaviour, we used scan data (N_scans_=27 068) to construct a multilevel multinomial behavioural model [44] with the Rstan package [45]. We linked one week of behavioural data (3 days before the faecal sample collection and 4 days after) to each faecal sample for the corresponding individual for both parasite data [11] and fGCM concentrations. We found no qualitative differences in estimates between the reduced and full focal datasets for the variables that could be included (results: S2).

Multilevel, multinomial behavioural models estimate the likelihood of a given behaviour from a set of categorical behaviours occurring at any given time in relation to a reference behaviour, while controlling for repeated observations from the same individual.

We set behaviour (feeding, resting, grooming given, grooming received, and moving) as our response variable, with moving as our reference variable. Moving was selected, as the reference variable is sensitive to frequency, and moving is a very common behaviour. We included parasite intensity (given as eggs per gram), parasite richness (number of genera), and NDVI as our primary fixed effects. We also controlled for other physiological effects by including fGCMs as a fixed effect, and we also controlled for sex, standardised rank and date. Individual ID and troop were included as random effects. In addition to summary statistics, we generated predicted probabilities for each behaviour for each predictor variable while holding other coefficients constant. This allowed us to look at changes in all behaviours, including the reference variable. Owing to the use of a reference behaviour (i.e., moving), coefficients of the multinomial model are not straightforward indicators of the effect of a predictor on the probability of performing a given behaviour [44] thus predicted probabilities are computed to understand the effects of the fixed effects on each behaviour.

## Model 2: The influence of parasite infection and ecology on behavioural predictability

We used entropy rate to determine whether parasite infection affects behavioural predictability. Entropy rate quantifies the predictability of the next observation, given the history of observations which occurred before it. Our entropy rate method estimates the distribution of behaviours (the frequency of each) and a transition matrix that describes transitions between behaviours [27]. An entropy rate of zero would indicate an individual engaged in a single behaviour for the entire observation period whereas an entropy rate of 1 indicates that an individual either engaged in multiple behaviours, switched behaviours frequently or both. As entropy rate has only been applied narrowly in animal behaviour, we began by validating its extension to observational data from wild monkeys, using both simulated and observed data (methods and results: supplementary S3). In order to estimate entropy rate, continuous focal samples were discretized into coded behavioural sequences. We assigned each behaviour a single letter code and created behavioural sequences by extracting behaviour from each focal at 30 second intervals, the optimal time period identified (N=693 faecal sample-matched sequences). We then used the ‘entropy’ package [46] in R version 3.5.2 (R Core Team, 2018), to calculate the entropy rate.

### Bayesian mixed-effects model structure

We constructed a mixed effects model with a Gaussian distribution in a Bayesian framework to assess the relationship between parasite intensity and behavioural entropy rate (distribution comparison results: supplementary material S4). Our response variable was behavioural entropy rate and, as with model 1, parasite intensity for ?*Protospirua* sp., *Trichostrongylus* sp., parasite richness and NDVI were included as our primary fixed effects while controlling for fGCM concentration, rank and sex as fixed effects. Given that individuals may be more likely to be active earlier in the morning and resting or grooming during the hottest part of the day, which may affect behavioural predictability, we included a spline on time of day as a fixed effect. Individual ID and troop were included as a random effects. As not all focal samples were exactly 10 minutes long, we also controlled for sequence length. We standardised continuous variables (rank, NDVI and sequence length) using one standard deviation (SDs) to allow comparisons of effect sizes across continuous and dichotomous variables. These variables were mean-centred on zero. We ran models with 4 chains and 2000 iterations which allows for a large enough sampling pool to achieve model convergence and conduct posterior sampling [47,48]. We used weakly informative priors (normal(0, 1)) and chain convergence was confirmed by 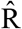 values ≤ 1.01. Model goodness-of-fit was assessed using the “posterior predictive check” (pp_check) function in the “bayesplot” package [49].

## Results

### Patterns of infection and co-infection

*?Protospirura* sp. had a mean annual sample prevalence of 98.74 % (±1.74 SD) and host group prevalence of 99.33% (±1.51 SD) with only 8/573 samples negative for all parasites. *Trichostrongylus* sp. had a mean annual sample prevalence of 22.04% (± 17.56 SD) and host group prevalence of 25.69% (±17.53 SD). Thus, all samples that were positive for Trichostrongylus sp., were also ?*Protospirura* sp. positive.

For *?Protospirura* sp., annual minimum and maximum egg counts from positive samples (ps) were 2 eggs per gram (EPG) and 5841 EPG respectively (mean_ps_ = 752.22, ±861.33 SD) while for *Trichostrongylus* sp., egg counts ranged from 2 to 47 EPG (mean_ps_ = 6.5, ±5.SD).

We found no evidence of a population-level relationship between ?*Protospirura* sp. infection intensity and *Trichostrongylus* sp. infection intensity (Estimate = 0.38, Estimate error = 0.64, lower 95% credible interval = −1.04, upper 95% credible interval = 1.56).

We found some evidence of inter-individual differences in random slopes for co-infection patterns of parasite intensity (Fig. 1). For some individuals, infection intensity of ?*Protospirura* sp. was high when *Trichostrongylus* sp. was absent or intensity is low. However, when *Trichostrongylus* sp. infection intensity was higher, ?*Protospirura* infection intensity was also high for some individuals. This pattern is stronger for some individuals than others. Note that estimate uncertainty is high for some individuals due to small individual-level sample size and this result should be interpreted with caution. Full model results are provided in the supplementary material (S5).

**Figure 1.**
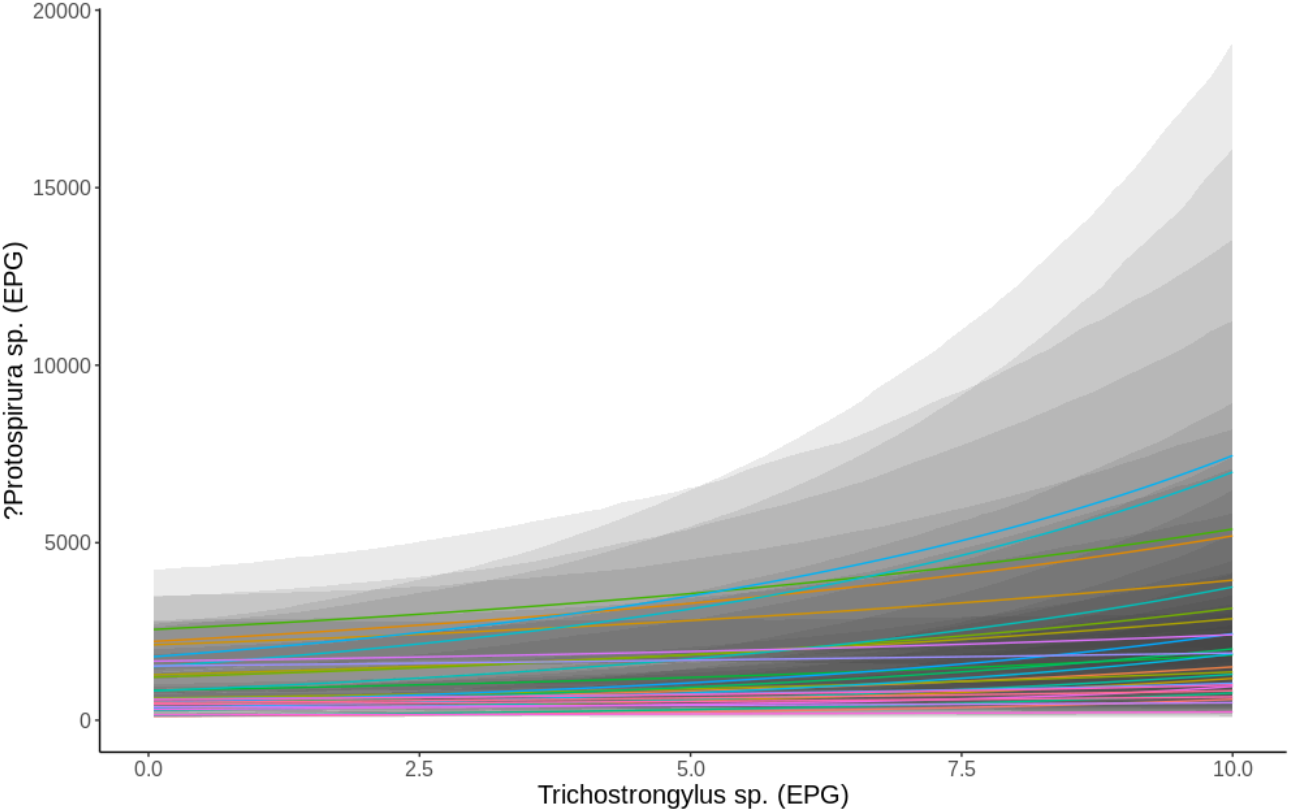
Estimate of faecal egg count of ?Protospirura sp. as a function of Trichostrongylus sp. faecal egg count derived from the fitted Bayesian mixed-effects model. Upper and lower 95% credible intervals (grey bands) shown.

### Model set 1: Influence of parasite infection and ecology on behaviour

#### Fixed effects

We found evidence of parasite-induced lethargy (i.e., increased resting time) and anorexia (i.e. reduced feeding time) as ?*Protospirura* sp. egg count increased (Fig. 2a). The probability of resting increased by 8.7% (l-CI = 2.2, u-CI =14.9) when egg counts were highest. This was predominantly traded off against moving, which showed a 7.4% (l-CI = 2.9, u-CI =12.2) and there was also a 4.3% decrease (l-CI = 0.16, u-CI =8.3) in the probability of foraging. The probability of both giving and receiving grooming were largely unchanged.

**Figure 2.**
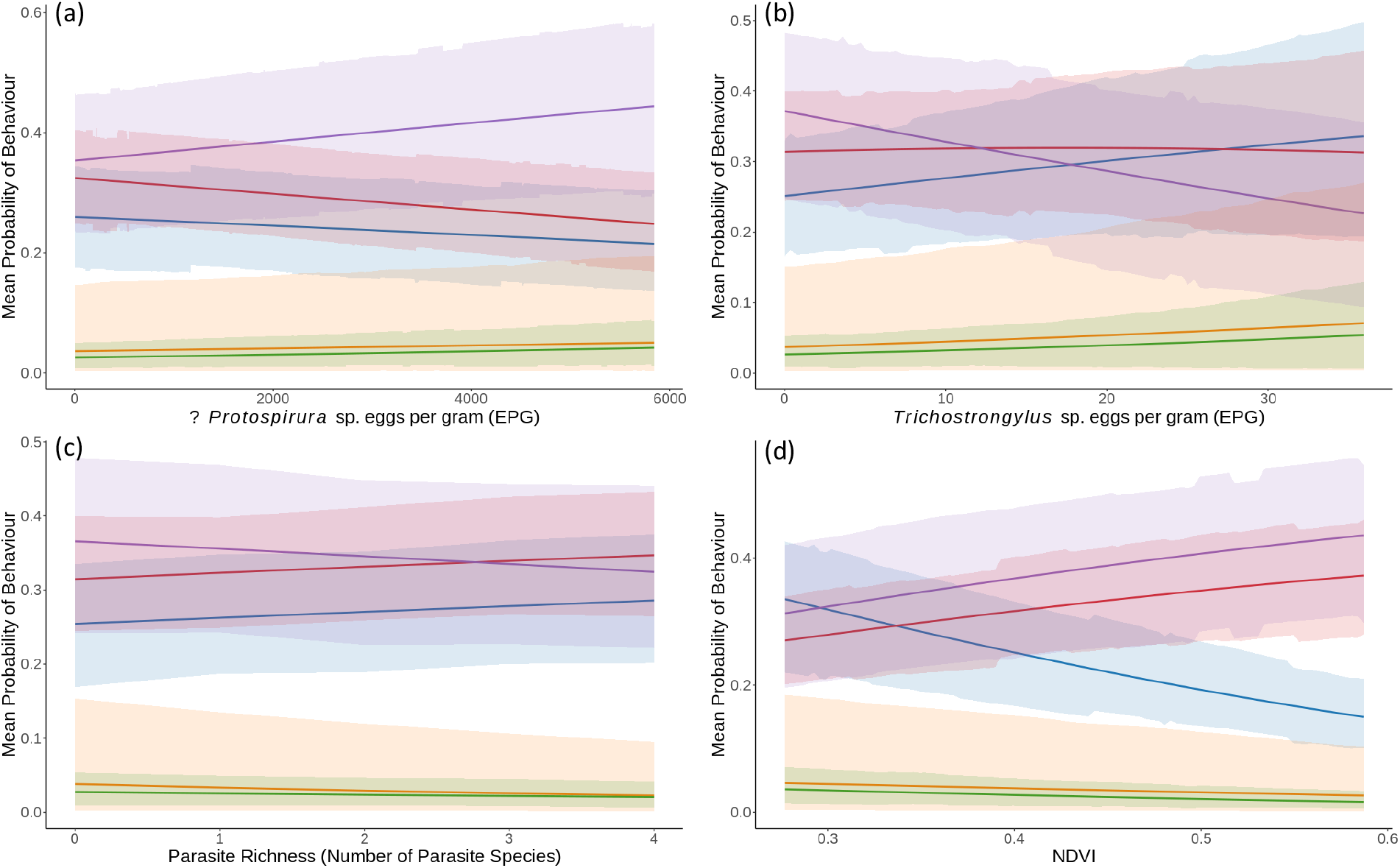
The relationships between the probabilities of behaviours being expressed as a function of each primary predictor variable. The 5 behaviours are: foraging (blue), resting (purple), moving (red), grooming in (green) and grooming out (orange). Shaded regions show 89% percentile intervals as calculated from the posterior samples

Conversely, we found that an increase in *Trichostrongylus* sp. loads resulted in a 15.4% (l-CI = 6.3, u-CI =24.6) reduction in the probability of resting. There was also an 8.8% (l-CI = 0.2, u-CI = 21.1) increase in the probability of foraging, while the probability of moving remained largely unchanged (Fig. 2b). The probability of both giving and receiving grooming increased slightly, by 4.0% (l-CI = −0.8, u-CI = 17.8) and 3.04% (l-CI = −1.6, u-CI = 11.4) respectively when *Trichostrongylus* sp. egg counts were higher; however, credible intervals were wide indicating uncertainty.

An increase in parasite species richness resulted in a slight decrease in the probability of resting (4.2%, l-CI = −1.7, u-CI = 10.4). However, credible intervals were wide and uncertainty high. Parasite richness did not influence the probability of the other behaviours occurring (Fig. 2c).

Although parasite intensity predicted changes in activity budget, the strongest predictor was change in food availability (Fig. 2d). When food availability was high, the probability of foraging decreased by 18.4% (l-CI = 12.3, u-CI = 23.8). This was accompanied by a 12.3% (l-CI = 8.1, u-CI = 16.0) increase in the probability of resting and a 10.1% (l-CI = 5.5, u-CI = 14.8) increase in the probability of moving. The probability of grooming given and received decreased slightly by 2.1% (l-CI = 0.09, u-CI = 7.9) and 1.9% (l-CI = 0.6, u-CI = 4.4), respectively. The full model output and summary can be found in the supplementary material (S6).

#### The influence of co-infection on behaviour

We found that, when *Trichostrongylus* sp. infection intensity was low (2 EPG), the probability of resting increased, feeding decreased and moving decreased as ?*Protospirura* sp. egg count increased (Fig 3). When *Trichostrongylus* sp. was high (35 EPG), the mean probability of resting was lower overall but still rose with increasing ?*Protospirura* sp. egg count and the probability of foraging decreased further. The probability of moving remained the same.

**Figure 3.**
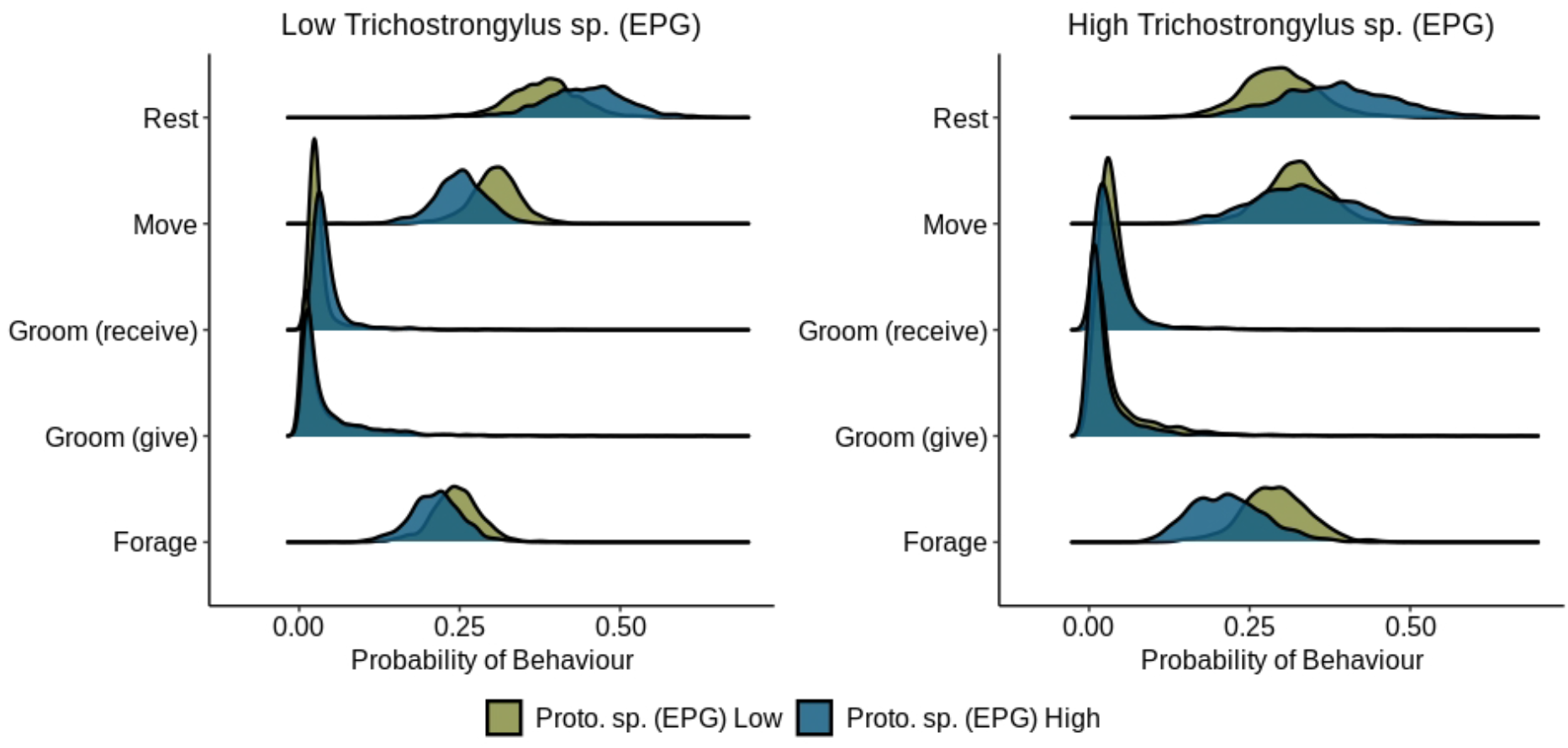
Changes in the mean probability of behaviours in response to high ?Protospirura sp. (Proto. sp.) when Trichostrongylus sp. intensity (EPG) was low (green) and high (blue). Density plots show probability of behaviours predicted by the model, with the height of the density curve indicating the probability of the predicted behaviour. The spread of the curve indicates the uncertainty.

### Model set 2: Influence of parasite infection and ecology on behavioural predictability

We found evidence of a positive relationship between NDVI and entropy rate (Est. = 0.10, Est. Error = 0.03, l-CI = 0.04, u-CI = 0.16). This indicates that an increase in food availability was associated with a decrease in behavioural predictability (Fig. 4). We found some evidence of a negative, non-linear relationship between entropy rate and time of day (sds Est. = 0.28, Est. Error = 0.23, l-CI = 0.13, u-CI = 0.87). Behavioural predictability was lowest in the early morning and increased until mid-day (Figure: supplementary S7).

**Figure 4.**
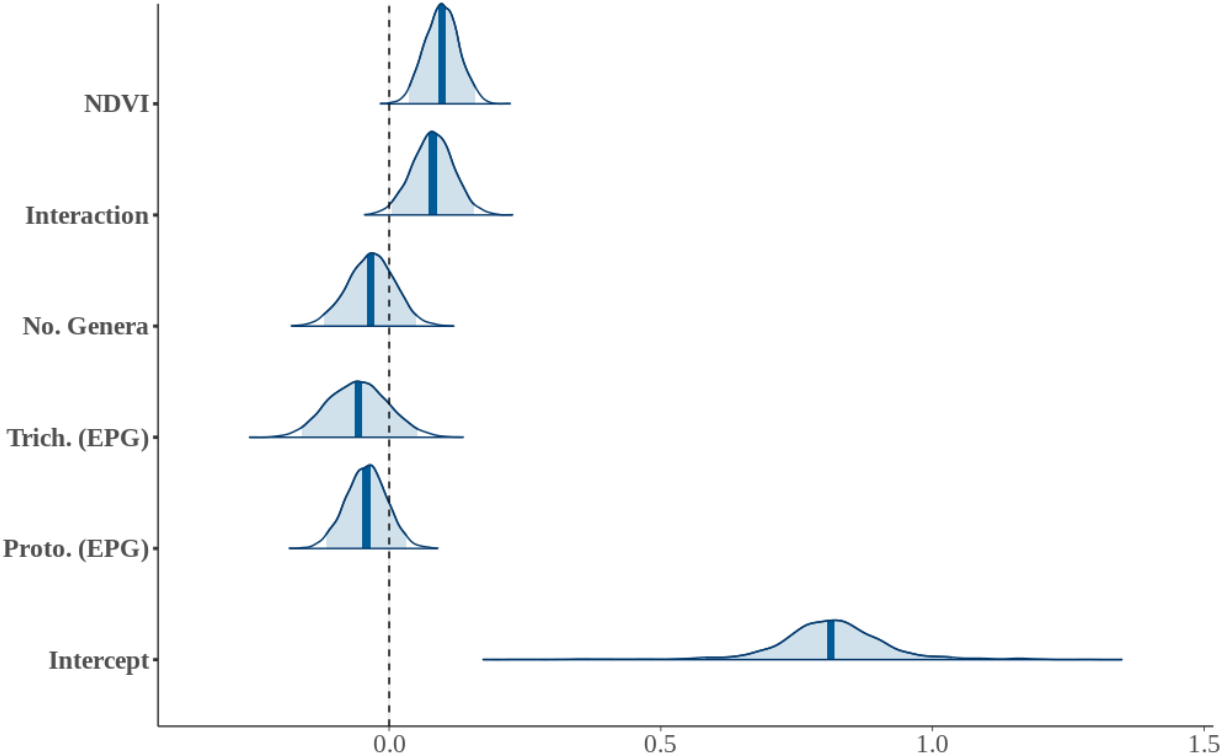
Posterior density plots from the GAMM showing the relationships between primary predictor variables and entropy rate. From top to bottom, variables are NDVI, interaction term between ?Protospirura sp. (EPG) and Trichostrongylus sp. (EPG), parasite richness (number of genera), Trichostrongylus sp. intensity (eggs per gram), ?Protospirura intensity (eggs per gram and the intercept. Vertical lines represent the mean and area under the curves show 95% credible intervals.

We found no evidence that ?*Protospirura* sp. and *Trichostrongylus* sp. parasite intensity or parasite richness influenced entropy rate (Fig. 4). Similarly, fGCM concentration, sex, rank and individual ID did not influence behavioural predictability. We found no effect of sequence length on entropy rate, which supports our use of a 6min focal cut off time. The full model only explained 9.2% of variance (R^2^ = 0.09, Est. Error = 0.02, l-CI = 0.06, u-CI = 0.13) suggesting there are other underlying drivers of behavioural predictability.

We found some evidence of a small, positive interaction between ?*Protospirura* sp. intensity (EPG) and *Trichostrongylus* sp. intensity. When *Trichostrongylus* sp. was low (2 EPG), entropy rate decreased with increasing ?*Protospirura* sp. intensity (Fig 5). Conversely, when *Trichostrongylus* sp. egg count was high, entropy rate increased with increasing ?*Protospirura* sp. infection intensity.

**Figure 5.**
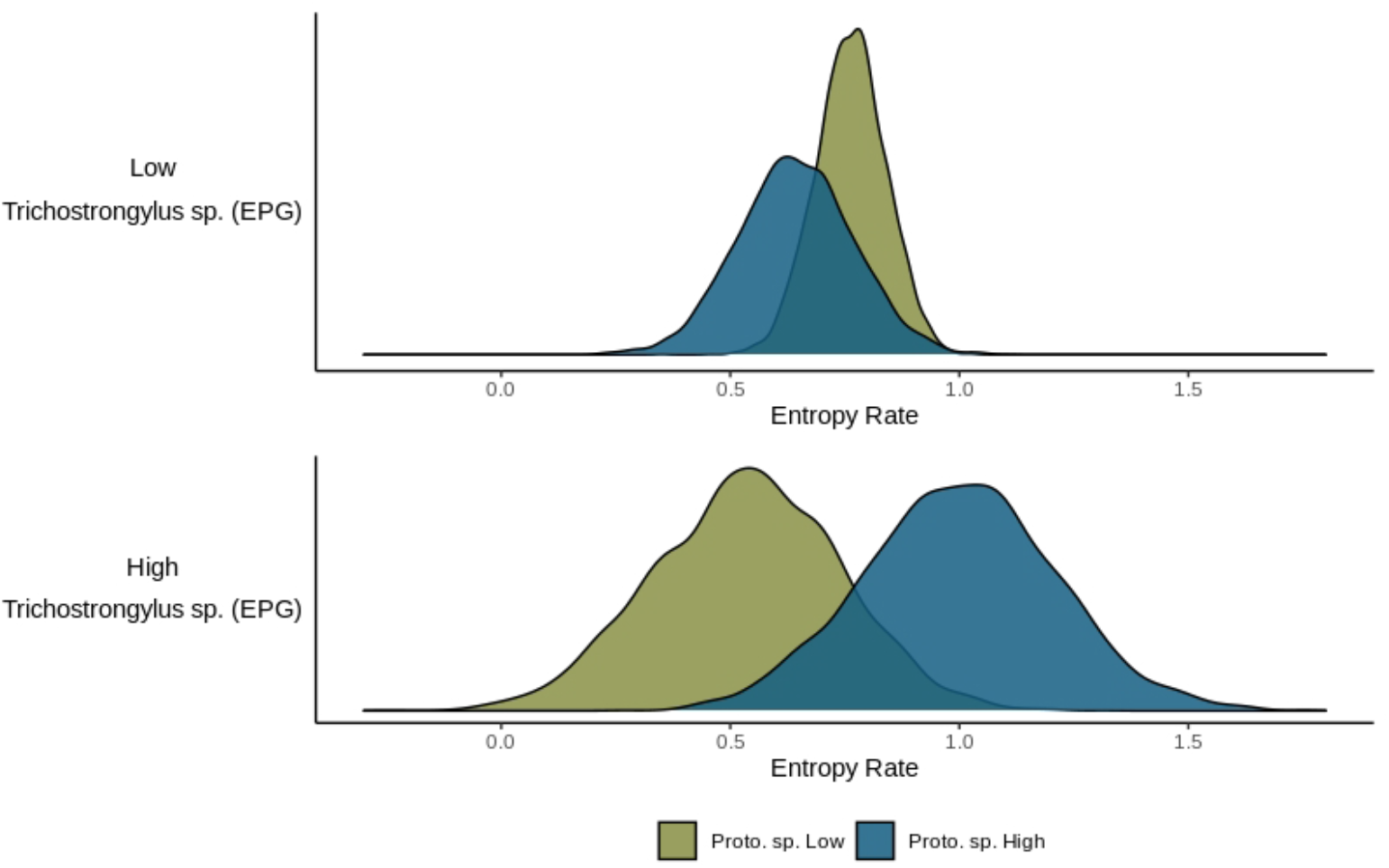
Changes in entropy rate in response to high ?Protospirura sp. (Proto. sp.) when Trichostrongylus sp. intensity (EPG) was low and high. Density plots show entropy rate predicted by the model, with the height of the density curve indicating the probability of the predicted entropy rate. The spread of the curve indicates the uncertainty.

## Discussion

Our results showed a relationship between parasite intensity and behavioural change, providing evidence for sickness behaviour in vervet monkeys. The nature of this relationship was not straightforward, however: we found that higher parasite loads predicted an increase in time spent resting, but that other behavioural changes were contingent on both the parasite species in question, and their interactions. This highlights the importance of considering multiple parasite infections when assessing the consequences of infection in wild non-human primates. Although we found evidence for changes in the overall amount of time devoted to particular activities, we found only limited evidence for more fine-grained changes in behavioural predictability (i.e., behavioural entropy rate) in response to parasite loadings. Given that food availability was the best overall predictor of behavioural change, it is likely that, for monkeys living in more extreme environments, coping with ecological stress overrides any fine-scaled ability to modulate behaviour in response to other stressors.

In line with previous work on non-human primates [1,11,18,20,50], we found evidence of sickness behaviour in response to two non-lethal gastrointestinal parasite infections. We found that increases in parasite intensity (EPG) of both ?*Protospirura* sp. and *Trichostrongylus* sp. resulted in changes in activity budget suggesting that these monkeys modify their behaviour in response to high parasite infection load. High ?*Protospirura* sp. parasite intensity resulted in “typical” sickness behaviour—increased resting, and reduced foraging and moving. This is notable as ?*Protospirura* sp. transmission relies on an intermediate arthropod host, so we might expect a positive relationship between foraging and increased parasite load. The inverse relationship in this case provides further support for the idea that what we see here is, indeed, sickness behaviour. It is possible that the change in behaviour is due to other underlying physiological processes that also occur when ?*Protospirura* sp. infection intensity is high. However, we found no relationship between faecal glucocorticoid metabolites (fGCM) concentration and behaviour, suggesting that changes in behaviour are a result of gastrointestinal parasite infection rather than an indication that individuals are coping with other stressors. Still, it is possible that this lack of relationship may also be a result of fGCM data collection not being fine-grained enough and a failure to detect more short-term increases in fGCMs. This emphasises the value of considering multiple physiological variables in assessing parasite-host relationships.

In the case of *Trichostrongylus* sp. we found a different pattern, where high infection intensity was associated with an increase in the amount of time spent foraging, along with a decrease in the probability of resting. The implication here is that different gastrointestinal parasites may exert different physiological pressures on the host and the manner in which they successfully cope with different non-lethal infections. For example, nutrition plays a vital role in a host’s ability to cope with the negative effects of gastrointestinal parasites [51], which could result in the need to forage more when *Trichostrongylus* sp. infection is high.

Alternatively, high *Trichostrongylus* sp. parasite intensity may coincide with other environmental or social changes that influence host behaviour or parasite dynamics. We found no relationship between temperature, rainfall, or NDVI and *Trichostrongylus* sp. parasite intensity (Blersch et al., in press.) suggesting that monkeys are not simply foraging more when *Trichostrongylus* sp. is high because food availability is lower.

We were also able to consider the co-occurrence of the two parasites. We found no strong relationship between *Protospirura* sp. and *Trichostrongylus* sp. faecal egg counts suggesting either that there is either a synergistic relationship, meaning one parasite species enhances the other, or that there is no direct competition between these two parasites [14]. We did find, however, that co-infection with these two nematodes resulted in different activity budget changes. When parasite intensity was high for both species, shifts in behaviour were different from those seen when only a single infection was considered. Specifically, we found that, when *Trichostrongylus* sp. infection intensity was high, monkeys still rested more with increasing ?*Protospirura* sp. egg count (i.e., showed the same pattern as when we considered ?*Protospirura* sp. infection alone), but they also moved more and decreased foraging further, which contrasts with the findings for ?*Protospirura* sp. alone. While the presence of both infections may also be linked to external environmental or social changes, it lends support to the theory that multiple infections exert differential changes on the host [14] and highlights the need to address co-infections when assessing animal health.

Contrary to some previous work on bats [9,52] and non-human primates [11], we found no marked change in the probability of either giving grooming or receiving grooming for individual infections, and only a small reduction in allogrooming when both ?*Protospirura* sp. and *Trichostrongylus* sp. infection intensity were high. While investment in sickness behaviour is fundamentally beneficial, and suppression of sickness behaviour can be detrimental to host fitness and survival, animals have to weigh the cost of modulating behaviours in response to infection [3]. Minimal change in grooming in response to infection intensity suggests these vervets maintain social relationships in the face of such external pressures. Young et al. [26], however, found that vervets engaged in fewer social behaviours when environmental conditions were sub-optimal. Given the harsh semi-arid environment, these vervets may be unable to further reduce the amount of time spent grooming in response to parasite infection; that is, they may have already reduced their grooming investment to the extent that any further reductions would incur unsustainable costs with respect to individual social benefits, and/or to group cohesion [6,7].

While our focus here was solely on time spent grooming, social interaction has been linked to infection susceptibility and transmission in several social species [53–55] including non-human primates [56,57]. This suggests that, despite the lack of change in the time spent grooming, increased parasite load may result in alternative suppressive strategies, such as changes in the number or identity of grooming partners. However, these strategies may be contingent on the route of parasite transmission which, for ?Protospirura specifically, is unlikely to be from direct transmission between individuals. More detailed grooming analysis is required to fully understand whether these vervets do, at least in part, modulate their grooming behaviour in response to infection and the risk that maintaining grooming frequency may incur. Alternatively, the relationship between grooming and parasite infection simply may be less clear given the lower time invested in grooming in comparison to other behaviours.

We also considered whether parasite infection intensity was linked to changes in behavioural structure. Behavioural entropy rate, derived from focal data, was not influenced by individual parasite infections but, when *Trichostrongylus* sp. infection intensity was high, entropy rate increased with increasing ?*Protospirura* sp. egg shedding. Thus, polyparasitism resulted in decreased behavioural predictability, indicating that monkeys engaged in more behaviours, changed behaviours more frequently, or both. This contrasts with studies on non-human primates that found a reduction in behavioural complexity or the rate of behavioural switching when individuals were parasite positive [11] or had impaired health [22,58]. Given that detrended fluctuation analysis [22,58] and the rate of behavioural switching [11] measure different aspects of behaviour, direct comparison between previous results and ours is difficult. However, our study shows that polyparasitism may be an important and more realistic consideration in the assessment of behavioural predictability or behaviour switching.

Although we found that parasite infections influenced both activity budgets and behavioural structure, the primary drivers of behavioural change were shifts in food availability; changes in both activity budget and behavioural structure were strongly linked to this. Previous work in our population has identified complex relationships between behaviour and environmental conditions, with food resources, temperature, rainfall, and standing water availability strongly influencing activity budgets and mortality [25,26]. Our findings here augment this previous work, providing the first evidence that food availability also affects behavioural structure: behavioural predictability decreased markedly when food availability was higher. This change resulted from a trade-off between a decrease in time spent foraging and an increase in both moving and resting when food availability was high. Changes in behavioural predictability have been shown to have short- and long-term consequences on fitness and survival, such as predator-prey interactions [59] and mating success [60]. However, beyond knowing that behavioural structure can serve as proxy measure of health [22], the implications for non-human primates are not yet well understood. Here, the use of entropy rate, rather than existing binary approaches, should allow us to identify the consequences of more complex behavioural trade-offs.

Taken together, our results provide the foundation for further research on both polyparasitism and the more fine-grained influences of non-lethal parasite infections on behaviour. We highlight the importance of using a detailed, comprehensive dataset when investigating how environment, physiology and parasitism interact to shape behaviour. Our findings also provide additional insight into how animals living in a harsh environment, with strong activity budget constraints, may adopt alternative approaches to parasite infection, avoidance, and transmission reduction.

## Acknowledgements

We thank the Tompkins family for permission to work at Samara Private Game Reserve and Kitty and Richard Viljoen for logistical support in the field. We are deeply grateful to all volunteer research assistants and students for their ongoing help with data collection, and thank Delaney Roth for assisting in focal sampling. We thank the Endocrine Research Laboratory for their help with the laboratory work at UP and Professor Cameron Goater (University of Lethbridge, Canada), who generously provided laboratory space as well as invaluable guidance during the analysis phase.

